# Interspike intervals within retinal spike bursts combinatorially encode multiple stimulus features

**DOI:** 10.1101/2020.02.13.947283

**Authors:** Toshiyuki Ishii, Toshihiko Hosoya

## Abstract

Neurons in various regions of the brain generate spike bursts. While the number of spikes within a burst has been shown to carry information, information coding by interspike intervals (ISIs) is less well understood. In particular, a burst with *k* spikes has *k*−1 intraburst ISIs, and these *k*−1 ISIs could theoretically encode *k*−1 independent values. In this study, we demonstrate that such combinatorial coding occurs for retinal bursts. By recording ganglion cell spikes from isolated salamander retinae, we found that intraburst ISIs encode oscillatory light sequences that are much faster than the light intensity modulation encoded by the number of spikes. When a burst has three spikes, the two intraburst ISIs combinatorially encode the amplitude and phase of the oscillatory sequence. Analysis of trial-to-trial variability suggested that intraburst ISIs are regulated by two independent mechanisms responding to orthogonal oscillatory components, one of which is common to bursts with different number of spikes. Therefore, the retina encodes multiple stimulus features by exploiting all degrees of freedom of burst spike patterns, i.e., the spike number and multiple intraburst ISIs.

**Author Summary:** Neurons in various regions of the brain generate spike bursts. Bursts are typically composed of a few spikes generated within dozens of milliseconds, and individual bursts are separated by much longer periods of silence (∼hundreds of milliseconds). Recent evidence indicates that the number of spikes in a burst, the interspike intervals (ISIs), and the overall duration of a burst, as well as the timing of burst onset, encode information. However, it remains unknown whether multiple ISIs within a single burst encode multiple independent information contents. Here we demonstrate that such combinatorial ISI coding occurs for spike bursts in the retina. We recorded ganglion cell spikes from isolated salamander retinae stimulated with computer-generated movies. Visual response analyses indicated that multiple ISIs within a single burst combinatorially encode the phase and amplitude of oscillatory light sequences, which are different from the stimulus feature encoded by the spike number. The result demonstrates that the retina encodes multiple stimulus features by exploiting all degrees of freedom of burst spike patterns, i.e., the spike number and multiple intraburst ISIs. Because synaptic transmission in the visual system is highly sensitive to ISIs, the combinatorial ISI coding must have a major impact on visual information processing.

## Introduction

Understanding the rules by which neuronal spike patterns encode information is essential for investigating the complex functioning of the nervous system [1, 2]. Neurons in various brain areas generate spike bursts, which are characterized by clusters of high-frequency spikes separated by longer periods of silence [3-5]. Burst spikes typically occur within the temporal window of postsynaptic integration (dozens of milliseconds), and thereby inducing synaptic response with higher probability than isolated single spikes [6-8]. In this regard, bursts are believed to represent an important neuronal code [7, 9, 10]. Previous analyses of burst information coding have suggested that the number of spikes within a burst [4, 11-19], the onset timing of a burst [5, 6, 20-22], and the duration of a burst [15, 23] all carry information.

Because a burst has multiple spikes, it has one or more intraburst interspike intervals (ISIs). In theory, these intraburst ISIs can carry information if, for example, they are modulated by sensory inputs. Such burst ISI coding should have significant effects on information transfer, because the efficiency of synaptic transmission is sensitive to ISIs [24]. Consistent with this idea, recent studies suggest that ISIs within bursts carry information [15, 19, 23, 25]. Although these studies have shown that the first ISI and average ISI within a burst carry information, interaction among multiple ISIs has been unclear. Theoretically, bursts with *k* spikes have *k*−1 intraburst ISIs, and these *k*−1 ISIs could encode *k*−1 independent values that represent information. Whether burst ISIs encode information in such a combinatorial manner is unknown.

In the vertebrate retina, retinal ganglion cells (i.e., the output neurons) generate spike bursts [3, 4, 26]. While the number of spikes within bursts encodes the amplitude of light intensity modulation [4], it is unknown whether intraburst ISIs encode information. In this study, using isolated salamander retinae, we investigated whether intraburst ISIs encode information regarding visual input. Our results indicated that intraburst ISIs encode oscillatory light intensity sequences different from the stimulus feature encoded by the spike number. When bursts contained three spikes, the two ISIs combinatorially encoded the amplitude and phase of the oscillatory components. Further analysis of the trial-to-trial variability suggested that intraburst ISIs are determined by two independent neuronal mechanisms that respond to two orthogonal oscillatory components. Collectively, our findings demonstrate that multiple ISIs within a retinal burst combinatorially encode multiple independent stimulus features that are different from that encoded by the spike number.

## Results

### Burst spike numbers encode the amplitude of light intensity modulation

We stimulated isolated larval salamander retinae using a spatially uniform visual stimulus with intensity modulation set at 30 Hz. Ganglion cell action potentials were recorded using a multi-electrode array. OFF ganglion cells, constituting the majority of the larval salamander retinal ganglion cells, generated spike bursts (Fig 1A). The majority of the spikes [82.0% ± 8.7%, mean ± SD (standard deviation), *n* = 41 cells] were observed in bursts comprised of two or more spikes, indicating that multi-spike bursts represent the major retinal code.

**Fig 1.**
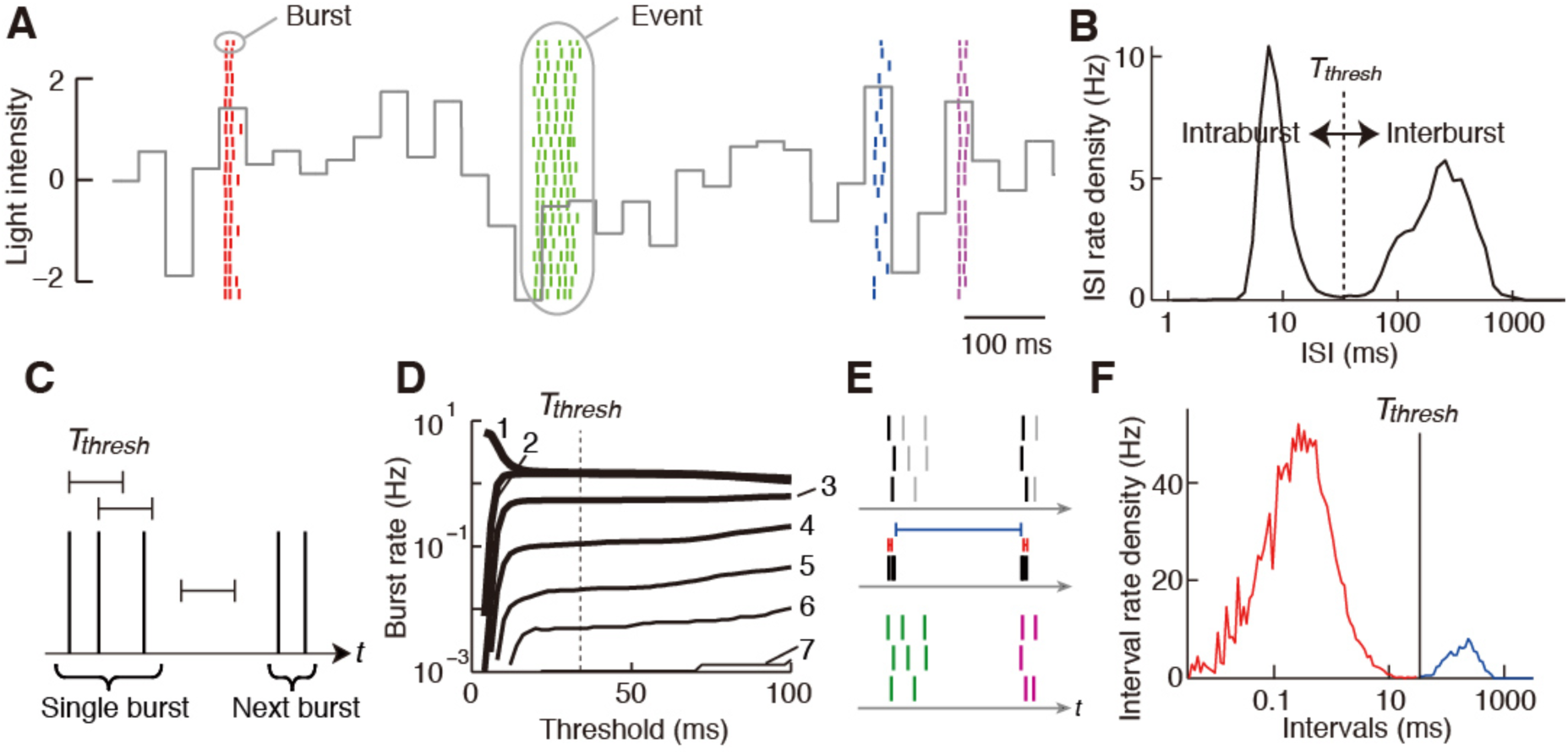
Retinal ganglion cells generate reproducible bursts. Data from a single salamander OFF ganglion cell. **(A)** Raster plot. Short vertical lines represent single spikes and each row shows spikes that occurred during a single repeat of the stimulation. Spikes of different events are shown in different colors. The gray continuous line shows the normalized light intensity of the stimulus (the mean and SD are 0 and 1, respectively). **(B)** ISI histogram. **(C)** Schematic illustration of the algorithm to define bursts. **(D)** Rates of isolated spikes (1) and bursts with 2–7 spikes (2–7) plotted against the threshold interval. **(E)** Schematic illustration of the algorithm to define events. **(F)** Histogram of intervals between merged onsets. The red and blue portions indicate intervals shorter and longer than *T*_thresh_, respectively.

During repeated presentation of the same stimulus, individual ganglion cells generated spike bursts at similar time-points across repeats (Fig 1A) [3, 4]. This reproducibility enabled the identification of corresponding bursts across repeats, which we termed “events” (Fig 1 and Materials and Methods) [3, 4]. Bursts generated in the same event had similar numbers of spikes in different repeats of the stimulus, while those in different events often had different numbers of spikes (Fig 1A). Accordingly, the number of spikes within bursts carried information about the stimulus (*p* < 0.01 for 41 of the 41 cells; the estimated mutual information was 0.80 ± 0.31 bits per burst, mean ± SD, *n* = 41). We next calculated the burst-triggered averages (BTAs), which represent the average stimulus sequence preceding isolated spikes and bursts with two, three, and four spikes (1-, 2-, 3-, and 4-BTA, respectively). The BTAs were sequences of different amplitudes (Fig 2A), and the difference between 1-BTA and 3-BTA had ON and OFF peaks around −170 and −40 ms relative to the burst onset (Fig 2B). This result indicates that the number of spikes within a burst encodes the amplitude of ON-to-OFF light intensity modulation within an interval of ∼130 ms.

**Fig 2.**
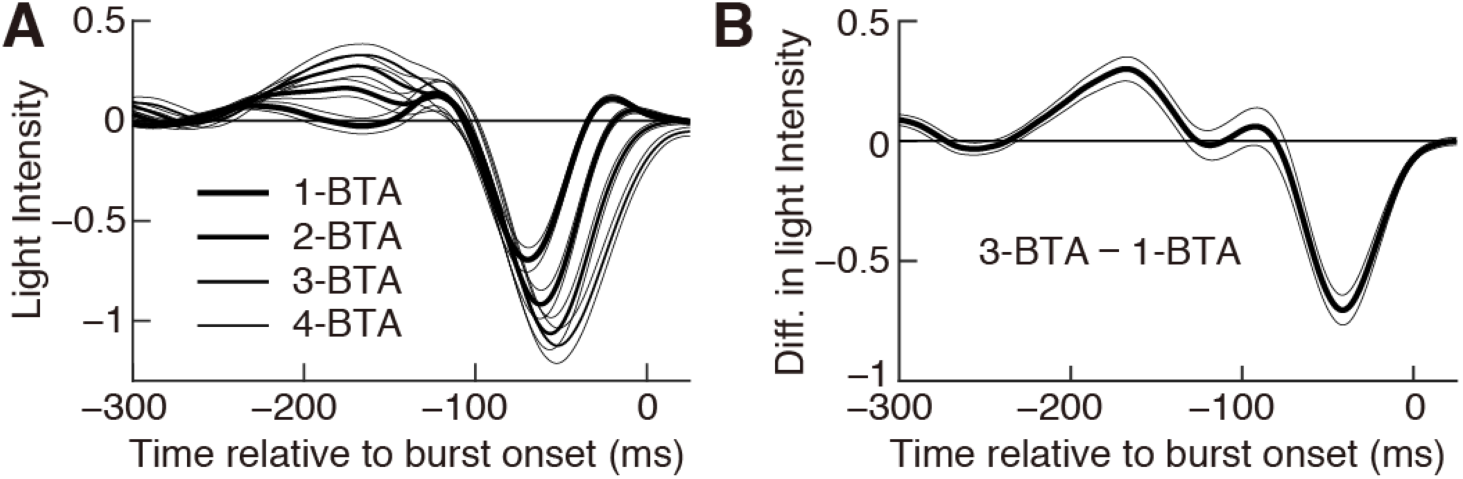
Spike number within a burst encodes the amplitude of light intensity modulation. (**A**) 1-, 2-, 3-, and 4-BTA indicate the averages of all stimulus sequences preceding isolated spikes, 2-, 3-, and 4-spike bursts, respectively. **(B)** Difference between 3-BTA and 1-BTA. Thick and thin lines indicate the average and standard error of mean (SEM) values, respectively. *n* = 41 cells.

### Burst ISIs encode oscillatory light intensity sequences

To investigate whether intraburst ISIs carry information, we first analyzed bursts composed of two spikes (2-spike bursts). For each ganglion cell, 2-spike bursts in the same event tended to have similar ISIs, whereas those in different events typically had different ISIs (Fig 3A). This suggested that intraburst ISIs convey information about the stimulus. Calculation of the mutual information confirmed this notion (*p* < 0.01 for 41 of the 41 cells; 0.33 ± 0.10 bits per burst, *n* = 41).

**Fig 3.**
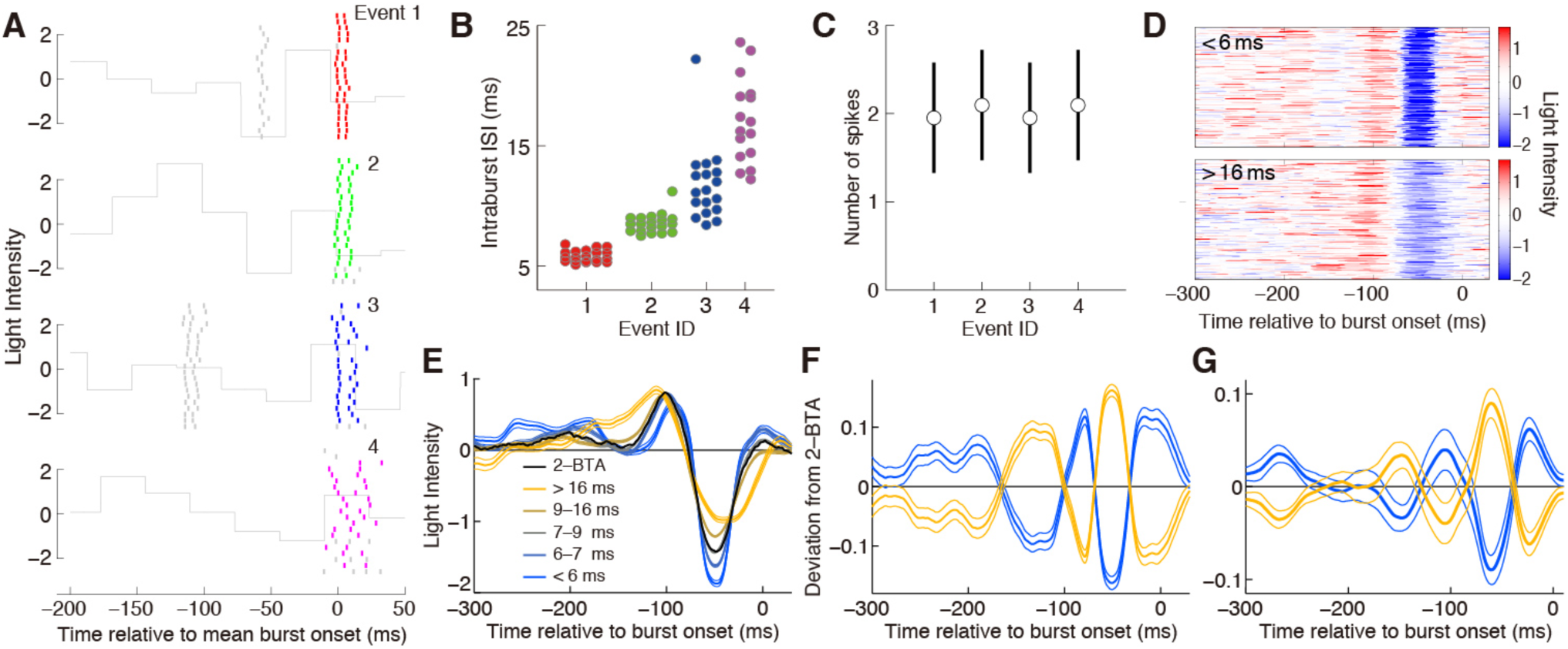
Intraburst ISIs of 2-spike bursts encode an oscillatory component of the visual input. **(A-F)** Data from the cell shown in Fig 1. **(A)** Raster plot. Colored lines represent 2-spike bursts. Short gray lines are the other spikes. The gray continuous line indicates light intensity, as shown in Fig 1A. Event IDs are shown. **(B and C)** Intraburst ISIs of 2-spike bursts (B) and the average and SD of the spike number (C) of the four events shown in (A). **(D)** Stimulus sequences preceding 2-spike bursts with an intraburst ISI of <6.0 (top) and >16.0 ms (bottom) are aligned with respect to the burst onset. **(E)** The thick black line indicates the average of all stimulus sequences preceding 2-spike bursts (2-BTA). Thick colored lines indicate the average of stimulus sequences preceding 2-spike bursts with different intraburst ISIs. Thin lines indicate SEM values. **(F)** Thick yellow and blue lines indicate the average of the stimulus sequence preceding 2-spike bursts with the longest and shortest 50% of intraburst ISIs, respectively, from which the 2-BTA is subtracted. The thin lines indicate SEM values. (**G)** Population analysis. Thick lines indicate the data shown by the thick lines in (F) averaged among 19 cells that generated at least 1500 2-spike bursts. Thin lines represent SEM values.

Two-spike burst ISIs were not correlated to the average number of spikes in events (Fig 3B and C; correlation = 0.0 ± 0.1, mean ± SD, *n* = 41), suggesting that these ISIs were modulated according to stimulus features different from those modulating the spike number. To characterize the stimulus features modulating 2-spike burst ISIs, we extracted the stimulus sequences preceding 2-spike bursts with different ISIs (Fig 3D and E). The results show that the stimulus sequences had systematic differences depending on the ISIs. We next subtracted the 2-BTA (black in Fig 3E) from the average of sequences preceding 2-spike bursts with long ISIs. The result was an oscillating sequence with two ON and two OFF peaks (yellow in Fig 3F and G). The subtraction from the 2-spike bursts with short ISIs gave the same sequence, but with the opposite sign (blue in Fig 3F and G). These results indicate that 2-spike burst ISIs encode an oscillatory deviation from the 2-BTA. Intervals between the ON and OFF peaks were ∼40 ms (Fig 3F and G) and, therefore, were much shorter than those of the stimulus feature encoded by the spike number (∼130 ms; Fig 2B). Thus, two-spike burst ISIs encode oscillatory sequences much faster than the intensity modulation encoded by the spike number.

### The two ISIs of three-spike bursts encode the phase and amplitude of oscillatory components

We next investigated the characteristics of 3-spike bursts. The first and second intraburst ISIs (ISI_1_ and ISI_2_) tended to be different for different events (Fig 4A and B) and carried information about the stimulus (*p* < 0.01 for 40 of the 41 cells; 0.46 ± 0.21 bits per burst, *n* = 41). In addition, the data suggest that ISI_1_ and ISI_2_ were modulated differently. For example, in Fig 4A, ISI_1_ was similar between the first (red) and second (green) events, but tended to be different for the third event (blue). In contrast, ISI_2_ was similar between the second and third events, but different for the first event. Accordingly, in the two-dimensional plot of ISI_1_ and ISI_2_, bursts of different events occupied different locations, and events did not align one-dimensionally, but were distributed two-dimensionally (Fig 4B).

**Fig 4.**
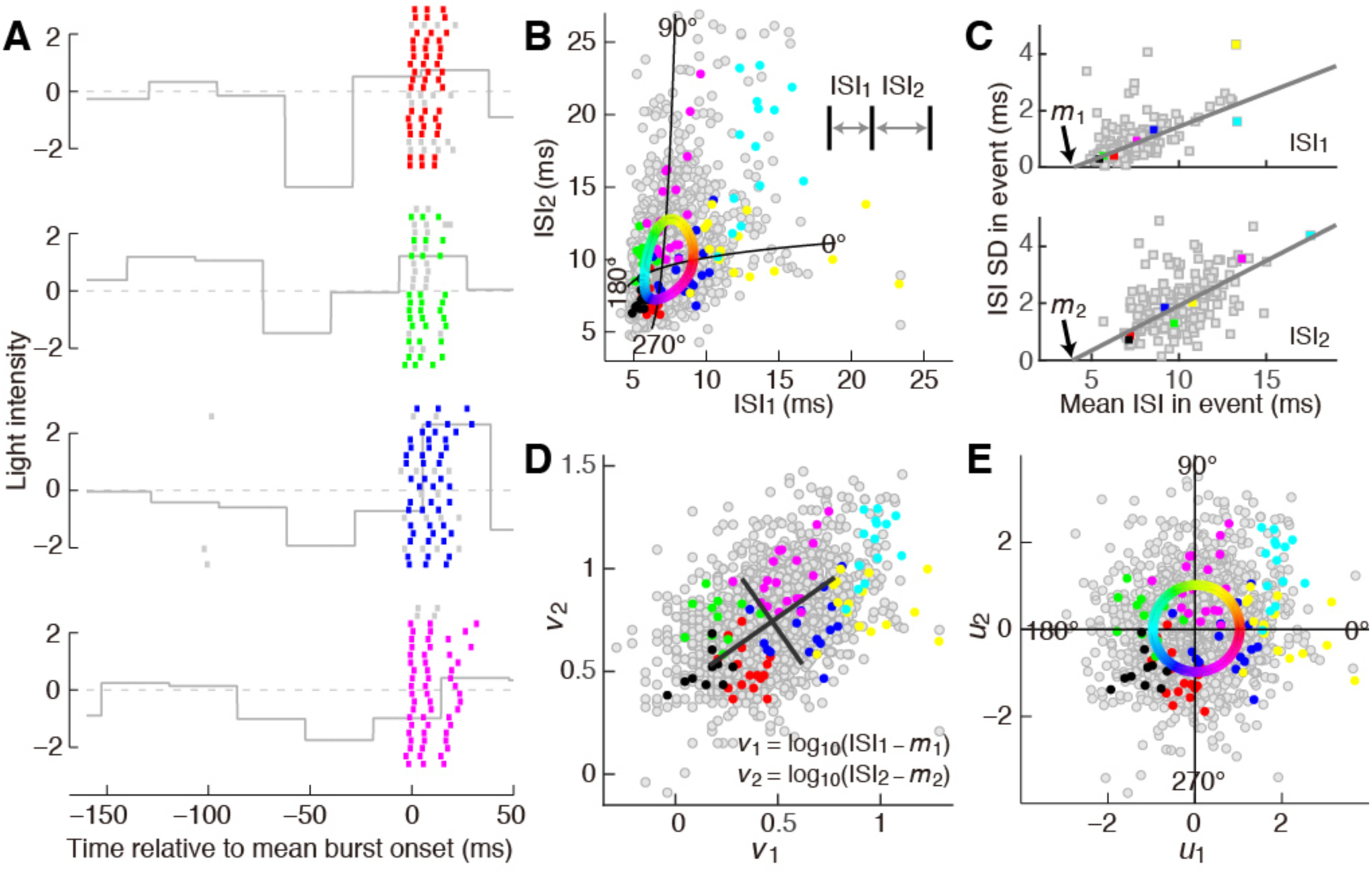
ISI patterns of three-spike bursts. Data from the cell shown in Fig 1. **(A)** Raster plot. Colored lines represent 3-spike bursts. Short gray lines are other spikes. The gray continuous line indicates the light intensity as shown in Fig 1A. **(B)** Distribution of ISI_1_ and ISI_2_. Each dot represents a 3-spike burst. Colored dots are bursts generated in seven different events, four of which are shown in (A) with the same color. Gray dots represent all other bursts. Black lines are the *u*_1_ and *u*_2_ axes. Burst phases are shown. The colors in the circle represent burst phases. **(C)** Variability of ISIs. Each dot represents an event. The horizontal and vertical axes are the average and SD of ISIs in an event, respectively. Gray lines show linear fits. **(D)** *v*_1_ and *v*_2_ were determined as indicated by the equations. Each dot represents a 3-spike burst. Black lines indicate the principle axes. The length of the lines represents the SD in the axis of the corresponding principle components. The coordinates of the crossing points are the averages of *v*_1_ and *v*_2_. **(E)** *u*_1_ and *u*_2_ determined by linear scaling of *v*_1_ and *v*_2_ along the principle axes. The colors in the circle show burst phases.

The above results suggest that the modulation of ISI_1_ and ISI_2_ has two degrees of freedom. The distribution of ISI_1_ and ISI_2_ had features that complicate further analysis. First, events with longer ISIs tended to have larger trial-to-trial ISI variability than events with shorter ISIs (Fig 4C). This inhomogeneous variability suggests that shorter ISIs represent information with a resolution higher than that represented by longer ISIs. Second, although ISI_1_ and ISI_2_ were modulated differently, they were not completely independent, but were correlated (Fig 4B; correlation coefficient = 0.16 ± 0.14, mean ± SD, *n* = 41). We were able to correct the first point by non-linearly scaling ISI_1_ and ISI_2_ so that different events had similar variability (Fig 4C and D). To correct the second point, variables were further linearly scaled to have an approximately circularly symmetric distribution so that the correlation was negligible (Fig 4E; correlation coefficient = −0.01 ± 0.13, *n* = 41). The resultant variables, *u*_1_ and *u*_2_, had a circularly symmetric distribution and, therefore, were approximately independent, with different events occupying a similar amount of area (Fig 4E).

Using *u*_1_ and *u*_2_, we defined the “burst phase” for each burst (Fig 4B and E). Stimulus sequences preceding 3-spike bursts exhibited systematic differences according to the burst phase (Fig 5A). We subtracted the 3-BTA (Fig 5B) from the sequences preceding 3-spike bursts. The resulting deviations were oscillatory components with two or three ON peaks separated by 70–80 ms, with OFF peaks among them (Fig 5C–E). The intervals between these peaks were almost constant, but their timing relative to the onset of bursts shifted depending on the burst phase (Fig 5C–E). When the burst phase increased from 0° to 360°, the peaks moved closer to the burst onset (Fig 5C–E), and the timing of the major ON peaks showed approximately linear dependence on the burst phase (Fig 5C–E). These results indicate that the phase of 3-spike bursts encodes the temporal phase of an oscillatory component. In addition, we found that the distance of the point (*u*_1_, *u*_2_) from the origin of the *u*_1_− *u*_2_ plane encodes the amplitude of the oscillatory component (Fig 5F). Similar coding was found for bursts elicited with natural scene movies (Fig 6).

**Fig 5.**
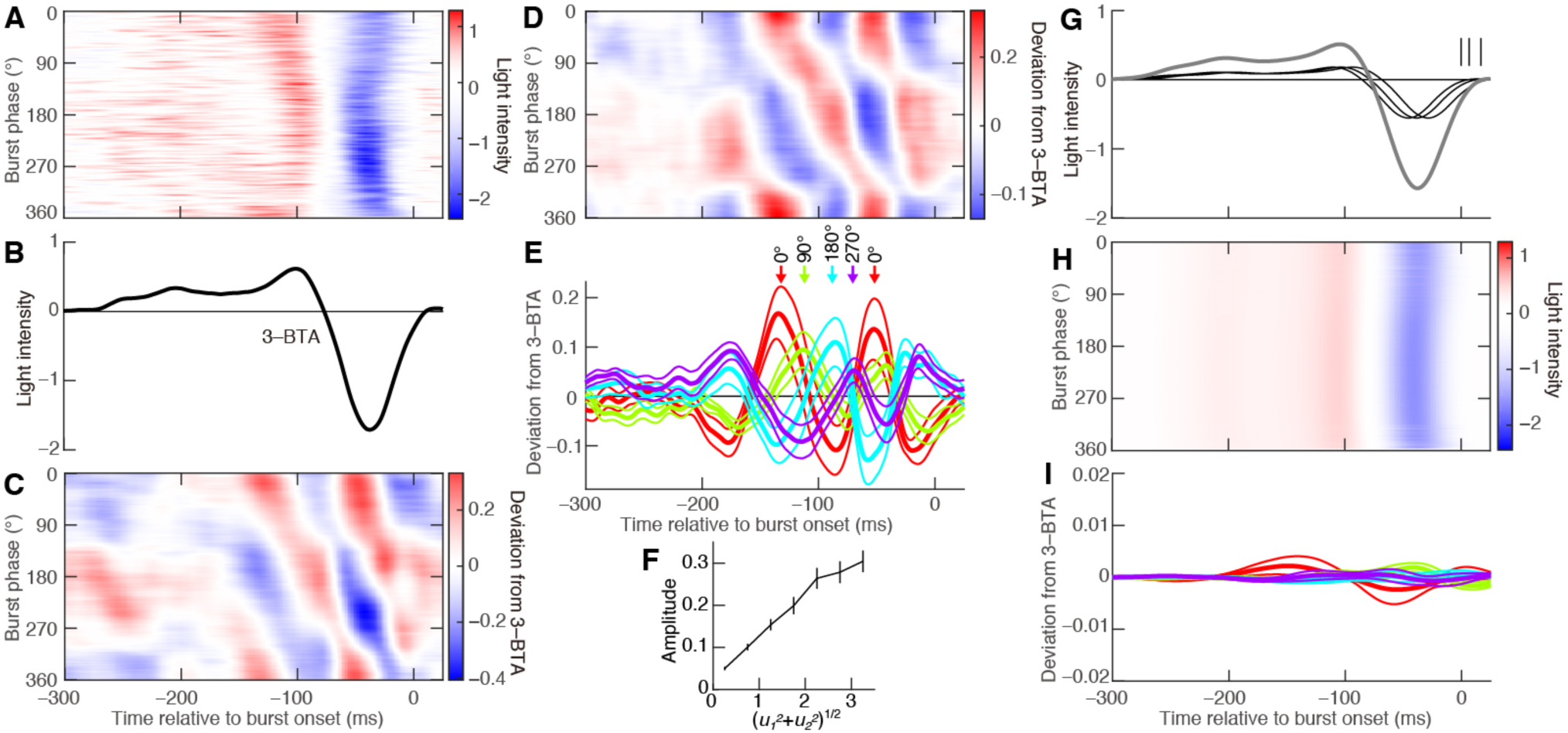
3-spike burst ISI patterns encode the phase and amplitude of oscillatory components. **(A–C)** Analysis of the cell shown in Fig 1. **(A)** Stimulus sequences preceding 3-spike bursts with different burst phases. **(B)** The average of all stimulus sequences preceding 3-spike bursts (3-BTA). **(C)** Data in (A) from which the 3-BTA is subtracted. **(D–F)** Population analyses. **(D)** Same analysis as in (C), averaged among 19 cells that generated at least 1200 3-spike bursts. **(E)** Thick lines indicate data in (D) at the indicated burst phases. Thin lines indicate the SEM values. Peaks around −100 ms are indicated. **(F)** Horizontal axis: 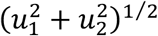, i.e., the distance of the point (*u*_1_, *u*_2_) from the origin in the *u*_1_–*u*_2_ plane in Fig 4E. Vertical axis: the root mean square of the oscillatory components between −200 and 25 ms. The error bars indicate SEM values for the 19 cells. **(G–I)** Reconstruction analyses. **(G)** Short vertical lines represent spikes in an example burst where ISI_1_ = 7 ms and ISI_2_ = 10 ms. Thin black lines indicate STAs calculated for 3-spike bursts of the cell used in (A–C), shifted according to the spikes in the example burst. Thick gray line indicates the reconstructed stimulus generated by adding the shifted STAs. **(H)** Analysis similar to (A) conducted for stimuli reconstructed as in (G) using the same bursts used in (A). Color-coding is as used in (A). **(I)** Population analysis similar to (E), conducted for reconstructed stimuli.

**Fig 6.**
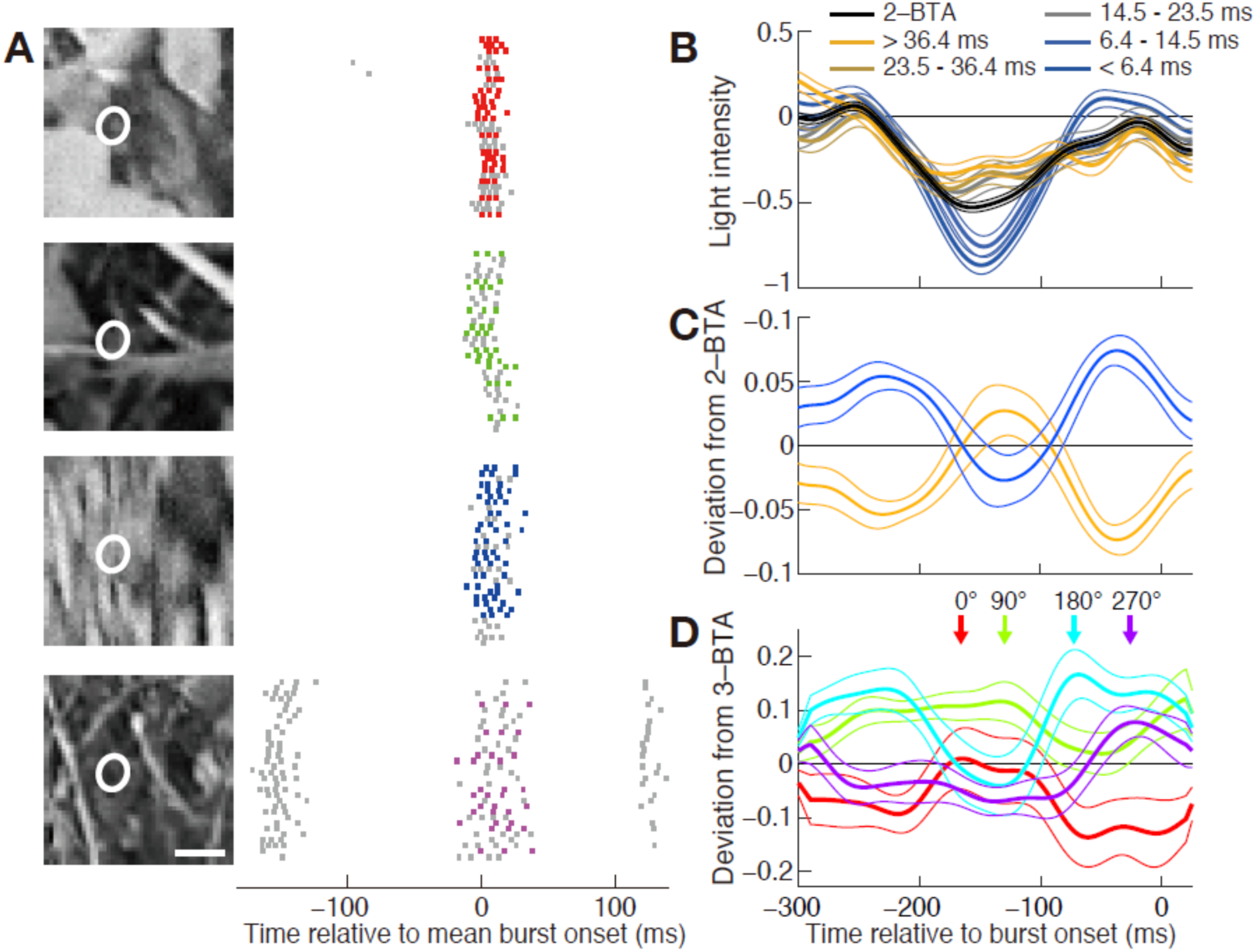
ISI analysis of bursts elicited by natural scene stimulation. **(A)** Responses of a single ganglion cell. (Right) Raster plots. The colored dots show 3-spike bursts. (Left) Image frames at −60 ms relative to the average timing of burst onsets. White ellipses show receptive field centers determined by reverse correlation and Gaussian fitting. Bar: 1 mm. **(B-D)** Analyses using light intensity at the receptive field center. **(B)** Analysis of 2-spike bursts of a single ganglion cell as shown in Fig 3E. Thin lines indicate SEM values. **(C)** Population analysis of 2-spike bursts similar to Fig 3G. Thick lines indicate the average of the stimulus sequence preceding 2-spike bursts with the longest (orange) and shortest (blue) 50% of intraburst ISIs, from which 2-BTA is subtracted. Thin lines indicate the SEM values. Data from 13 cells that generated more than 800 2-spike bursts. **(D)** Population analysis of 3-spike bursts similar to Fig 5E. Stimulus sequences preceding 3-spike bursts with different phases. The 3-BTA is subtracted. The thick and thin lines indicate the average and SEM values among 8 cells that generated more than 1000 3-spike bursts.

As shown in Fig 4B, bursts with the phase 0° and 180° had only ∼5- and ∼1-ms differences in ISI_1_ and ISI_2_, respectively. Nevertheless, for bursts with the phase 0°, the ON and OFF peaks in the encoded sequences were separated by ∼80 ms, while the separation was only ∼50 ms for bursts with the phase 180° (Fig 5A). These results suggest that single spikes in bursts do not simply indicate the occurrence of a single characteristic in light intensity sequence. To further confirm this point, we conducted a simple reconstruction analysis. We calculated the spike-triggered averages (STA) for three-spike bursts and then generated an estimated stimulus sequence by adding the STA aligned according to the burst spikes (Fig 5G). This reconstruction failed to reproduce the observed dependence of the ON and OFF peaks on the burst phases (compare Fig 5A and H), and the deviation from the 3-BTA was much smaller than that of the actual data (compare Fig 5E and I). To quantify the amplitude of the deviation, the root mean square of the deviation from the 3-BTA was calculated for the period between −200 and +25 ms and averaged for all burst phases. The values were significantly larger for the actual stimuli than for the reconstructed stimuli (0.107 ± 0.061 for actual stimuli and 0.003 ± 0.002 for reconstructed stimuli, *n* = 19, *P* = 1.5 × 10^−7^, two-tailed Mann–Whitney–Wilcoxon test). Thus, the simple reconstruction model failed to explain the observed burst coding.

### Two independent components of burst patterns

The oscillatory component encoded by 2-spike burst ISIs had peak-to-peak intervals that are similar to those of the components encoded by 3-spike bursts (∼80 ms; compare Fig 3G and 5E), suggesting that 2- and 3-spike burst ISIs are modulated by related stimulus features. To further characterize this similarity, we analyzed events in which both 2- and 3-spike bursts were generated (e.g., Fig 4A, bottom). For each of these events we calculated the average values of *u*_1_ and *u*_2_ for 3-spikes bursts and the average value of the 2-spike burst ISIs (Fig 7A). Plotting the data on the *u*_1_-*u*_2_ plane showed that 2-spike burst ISIs differ systematically depending on the position of the events along the direction of ∼30° (Fig 7B). A linear fitting indicated that the optimum direction was 33.1° ± 15.6° (circular mean ± SD, *n* = 41; Fig 7C, magenta), suggesting that 2-spike burst ISIs are modulated by an oscillatory component that modulates 3-spike burst patterns in the orientation of ∼33.1° on the *u*_1_-*u*_2_ plane.

**Fig 7.**
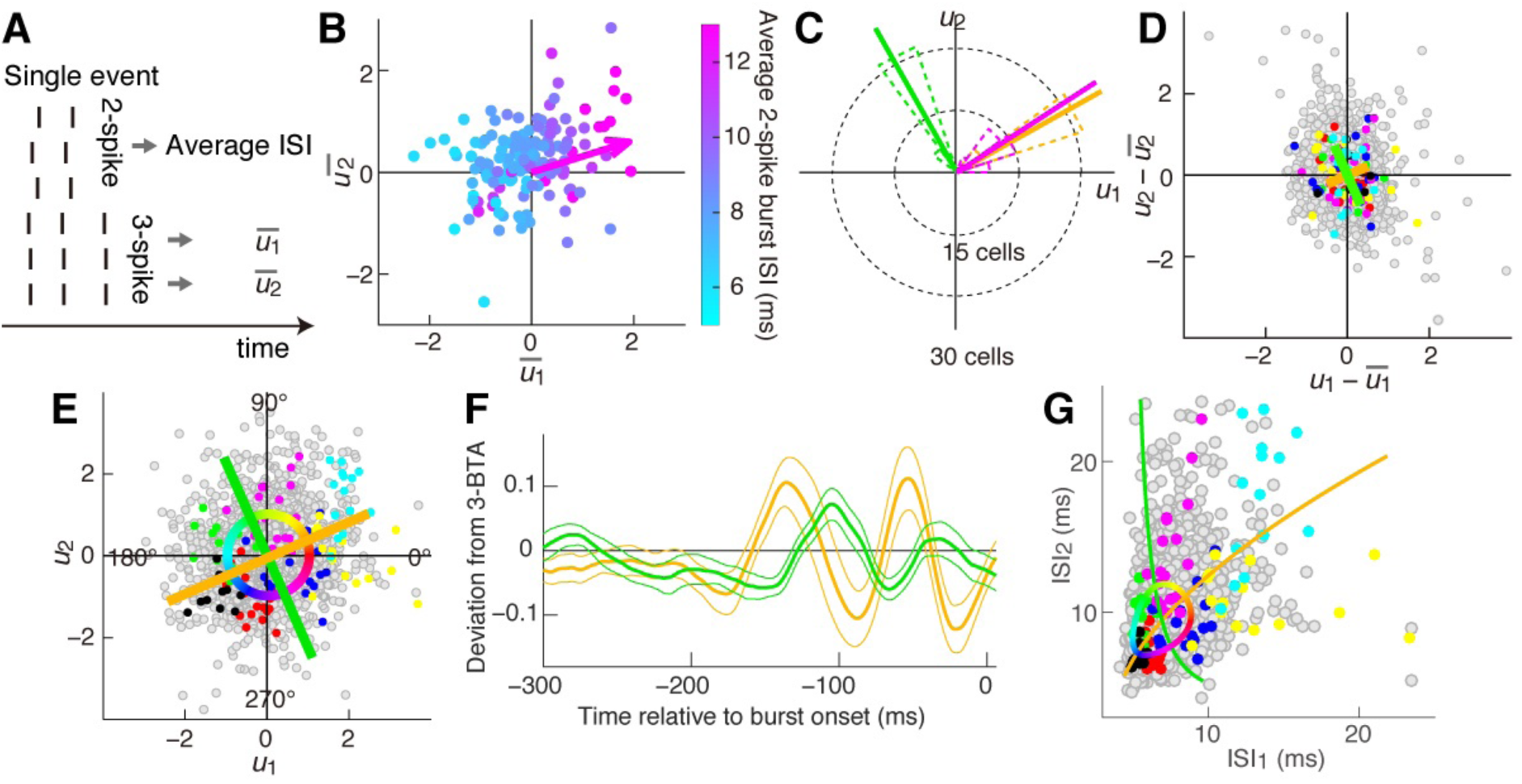
Identification of independent components that determine burst patterns. **(A)** The average value of 2-spike burst ISI and the average values of 3-spike burst *u*_1_ and *u*_2_ were determined for each event. **(B)** Dependence of 2-spike burst ISIs on 3-spike burst patterns. Each dot represents an event in which the cell generated both 2- and 3-spike bursts. The horizontal and vertical axes are the averages of *u*_1_ and *u*_2_ of 3-spike bursts in an event, respectively. The color indicates the average ISI of 2-spike bursts. The arrow shows the orientation of the increase of 2-spike burst ISIs determined by linear fitting. **(C)** Population analyses for *n* = 41 cells. (Magenta) The orientation of the increase of 2-spike burst ISIs determined as in (B). (Orange and green) Principle axes of trial-to-trial variations of *u*_1_ and *u*_2_, determined as in (D). Orange shows the shorter axes. Dotted and thick lines are the circular histograms and average orientations, respectively. **(D–F)** Analyses of trial-to-trial variability. **(D)** Trial-to-trial variations of *u*_1_ and *u*_2_ in each event. Each dot represents a 3-spike burst. The orange and green lines indicate the orientation of the principle components through all events. The lengths represent the SD along the axes. Compare with Fig 4E. **(E)** The same panel as Fig 4E, shown with the orientations of the principal components in (D). **(F)** Stimulus sequences preceding bursts with the phases corresponding to the two components, as shown in Fig 5E. Data from 19 cells that generated at least 1200 3-spike bursts. **(G)** The axes and circle in (E) are plotted on the ISI_1_–ISI_2_ plane.

The above result raises the hypothesis that the retinal mechanism that modulates 2-spike ISIs also modulates 3-spike burst patterns in the orientation of ∼33.1° on the *u*_1_-*u*_2_ plane. Since *u*_1_ and *u*_2_ are suggested to have two degrees of freedom, one possibility is that another independent mechanism modulates 3-spike burst patterns in the orthogonal orientation, i.e., ∼123.1°, on the *u*_1_-*u*_2_ plane. If two such independent mechanisms were present, modulation of *u*_1_ and *u*_2_ in the two orthogonal orientations would have independent trial-to-trial variations, and we tested this prediction. Although *u*_1_ and *u*_2_ had an approximately circularly symmetric distribution (Fig 4E), their trial-to-trial variations within each event had an asymmetric distribution (Fig 7D). Principal component analysis of this distribution indicated that the principal axes corresponding to the smaller and larger variances were in the orientations of 29.4° ± 7.3° and 119.4° ± 7.3° (*n* = 41; orange and green in Fig 7C and D). This analysis indicates that modulations of *u*_1_ and *u*_2_ in these two orientations have approximately independent trial-to-trial variation. Because these axes (∼29.4° and ∼119.4°) are close to those proposed for the hypothesis (∼33.1° and ∼123.1°, see Fig 7C), the results are in accordance with the presence of two independent mechanisms, one of which (∼30°) is common to 2- and 3-spike bursts. Consistently, the oscillatory component encoded by 3-spike bursts with the phase of the common component orientation (29.4° ± 7.3°) was similar to that encoded by 2-spike burst ISIs (compare Fig. 7F, yellow and Fig 3G, yellow). Given that the total distribution of *u*_1_ and *u*_2_ is circularly symmetric, the smaller trial-to-trial variability of the ∼30° component as compared with the ∼120° component indicates that the former is more precise. Therefore, the component common to 2- and 3-spike bursts is more informative than that specific to 3-spike bursts.

The two approximately independent components encode oscillatory components that are approximately orthogonal to each other, i.e., components with ∼1/4 cycles difference in phase (Fig 7F). This result suggests that the two orthogonal oscillatory stimulus components modulate 3-spike burst patterns in the orthogonal orientations on the *u*_1_-*u*_2_ plane. This model is consistent with the above result that 3-spike burst patterns encode the amplitude and phase of an oscillatory component, since an oscillatory sequence with an arbitrary amplitude and phase can be approximated by a sum of two orthogonal oscillatory components with fixed phases and the same frequency.

## Discussion

Our results reveal that intraburst ISIs of retinal bursts encode oscillatory light intensity sequences that are much faster than the sequence encoded by the spike number. When a burst has three spikes, the two intraburst ISIs combinatorially encode the amplitude and phase of the oscillatory component. These results therefore suggest that a *k*-spike burst (*k* = 2, 3) encodes *k* different stimulus features by exploiting all the *k* degrees of freedom, i.e., the spike number and *k*−1 ISIs. This simultaneous representation of multiple stimulus features enables multiplexed information coding, a mechanism that greatly increases the information transmission capacity [19, 27, 28]. Whether this combinatorial coding occurs for bursts with *k* ≥ 4 spikes remains unknown, as these bursts were rarely observed under our experimental conditions (Fig 1D).

### Mechanisms of the combinatorial ISI coding

Figure 8 shows a coding model that is consistent with our findings. The amplitude of slow light intensity modulation determines the spike number within a burst. Intraburst ISIs are regulated by two independent mechanisms that are driven by orthogonal fast oscillatory stimulus components, as suggested by the comparison between 2- and 3-spike bursts and the analysis of trial-to-trial variation. When a burst contains two spikes, the ISI is regulated by one of the two mechanisms, and thus 2-spike burst ISIs encode the amplitude of an oscillatory component of a fixed phase. When a burst has three spikes, the two mechanisms combinatorially determine the two ISIs. Because the two mechanisms are driven by the two orthogonal oscillatory components, the two ISIs of 3-spike bursts carry information about both the amplitude and phase of the oscillatory component. Modulation of the 3-spike ISI pattern by the common component is similar to modulation of the burst duration, i.e, ISI_1_ + ISI_2_ (see yellow in Fig 7G).

**Fig 8.**
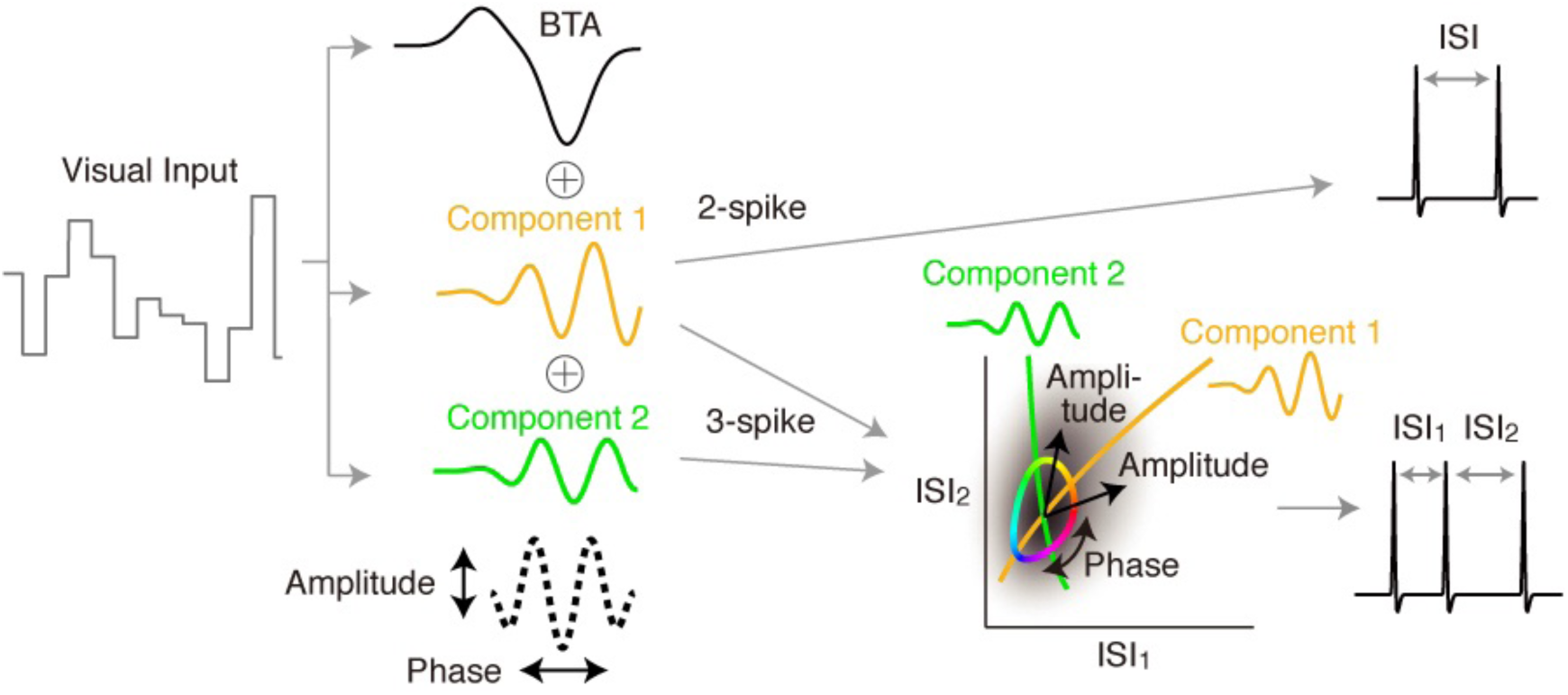
Schematic view of burst coding. The dotted line indicates the sum of the oscillatory components 1 and 2, whose amplitude and phase are encoded by the 3-spike burst pattern.

The two proposed mechanisms modulating intraburst ISIs exhibit approximately independent trial-to-trial variation, raising the possibility that the two mechanisms rely on two largely non-overlapping synaptic pathways. Such circuits, if present, may have different temporal properties, considering that the two mechanisms respond to two different temporal sequences. In the vertebrate retinae, bipolar cells have ∼10 subtypes [29-32], and different subtypes have distinct physiological properties [32-34] and varying temporal response characteristics [35-37]. In addition, the inhibitory effect of amacrine cells on bipolar cells generates further variation of temporal properties [38]. Therefore, specific subsets of bipolar and amacrine cells may constitute largely non-overlapping synaptic pathways underling the combinatorial coding.

### Implications for visual processing

It is currently unclear to what extent the burst ISI information analyzed in this study is transmitted to the brain. However, in many brain regions, neuronal responses are sensitive to the millisecond-scale temporal structure of synaptic inputs [24]. For example, in synaptic transmission from the retina to the lateral geniculate nucleus (LGN), retinal spikes with ISIs of a few milliseconds are much more effective in eliciting LGN spikes than those with ISIs of > ∼20 ms [39-45]. Consistently, LGN burst ISIs are sensitive to the millisecond-scale structure of current input [19]. Similar to synaptic connections from the retina to the LGN, those from the LGN to cortical neurons are more responsive to spikes with short ISIs than those with long ISIs [46]. Such dependence of neuronal responses on input ISIs suggests that bursts with different ISIs elicit different spike responses of postsynaptic neurons. In addition, the dependence on ISIs varies among individual synaptic connections [44-46]. This variation suggests that individual synapses have different preference for bursts with different ISIs and thus may function as a system to decode burst ISIs [24]. Although the present study investigated the dependence of ISIs on the temporal patterns of visual stimuli, retinal ISIs also depend on the spatial patterns [47]. Therefore, it is possible that burst ISIs encode spatial information as well as temporal information.

### Conclusions

The present results suggest that the retina employs mechanisms to regulate multiple components of intraburst ISIs, and thereby encodes multiple stimulus features by exploiting all degrees of freedom of burst spike patterns, i.e., the spike number and multiple intraburst ISIs. This burst coding is likely to affect visual information transmission, as synaptic transmission is sensitive to ISIs. Because bursts occur in various regions of the brain, analyses similar to the present study may reveal previously overlooked information transmission in those regions.

## Materials and Methods

### Animals

All experiments were approved by the RIKEN Wako animal experiments committee and were performed according to the guidelines of the animal facilities of the RIKEN Center for Brain Science. Larval tiger salamanders were provided by Charles D. Sullivan Co. Inc., Nashville, Tennessee, USA.

### Recording and Stimulation

Retinal recording was performed as described previously [48]. Dark-adapted retinae from larval male and female tiger salamanders were isolated in oxygenated Ringer’s medium at 25 °C. A piece of the retina (2–4 mm in width) was mounted on a flat array of 61 microelectrodes (MED-P2H07A, Alpha MED Scientific Inc., Ibaraki, Osaka, Japan) and perfused with oxygenated Ringer’s solution (2 mL/min; 25 °C). Spatially uniform white light (intensity refreshment at 30 Hz; mean and SD of the intensity were 4.0 and 1.4 mW/m^2^, respectively) was projected through an objective lens using a CRT monitor (60-Hz refresh rate; E551, Dell Inc., Round Rock, Texas, USA) controlled by the Matlab Psychophysics Toolbox [49, 50] or a light-emitting diode (E1L53-AWOC2-01 5-B5, Toyoda Gosei, Japan). The light intensity sequence was a random Gaussian sequence (65.5–183.3 s). The same sequence was repeated typically more than 20 times. For the natural scene stimulation, 200 s of a movie [51] was projected at 30 Hz using the CRT monitor (64 × 64 pixels, 60.6 μm/pixel; the mean intensity was 4.0 mW/m^2^). Amplified voltage signals from the electrodes were stored and action potentials of single units were isolated using a Matlab program (a gift from Dr. Stephan A. Baccus). Analyses were performed using stable cells with mean firing rates >1.7 Hz.

### Identification of Bursts and Events

Histograms of the ISIs were generated for each isolated ganglion cell. The histograms often had two distinct peaks (Fig 1B) representing shorter and longer ISIs, corresponding to intra- and interburst ISIs, respectively [3-5]. The threshold interval *T*_*thresh*_ was set at the trough between the two peaks in the ISI histogram (Fig 1B). *T*_*thresh*_ was 38.6 ± 20.0 ms (mean ± SD) for 41 cells stimulated with the spatially uniform stimulation, and 87.7 ± 18.5 ms for the 16 cells stimulated with the natural scene movie. If two consecutive spikes occurred with an interval shorter than *T*_*thresh*_, they were incorporated into the same burst, while they were separated into two consecutive bursts if the interval was longer than *T*_*thresh*_ (Fig 1C). The robustness of this method was examined as follows. Bursts were defined using various threshold intervals of ∼10 ms to ∼100 ms, and the rates of the isolated spikes and bursts with 2–7 spikes were measured (Fig 1D). *r*_−10_, *r*_0_, and *r*_+10_ (Hz) denote the rates of the 2-spike bursts defined by the threshold intervals *T*_*thresh*_−10 ms, *T*_*thresh*_, and *T*_*thresh*_+10 ms, respectively. The maximum rate change, max(|*r*_−10_ − *r*_0_|, |*r*_+10_ − *r*_0_|)/*r*_0_, was 0.021 ± 0.020 (mean ± SD, *n* = 41) for cells stimulated with the spatially uniform stimulation, and 0.022 ± 0.014 for the 16 cells stimulated with the natural scene movie. These small values indicate the robustness of the method. The median intraburst ISIs of the 2-spike bursts and the median of the duration (ISI_1_ + ISI_2_) of the 3-spike bursts were 8.1 ± 2.8 and 14.4 ± 4.7 ms (mean ± SD, *n* = 41), respectively, for the cells stimulated with the spatially uniform stimulation, and 15.4 ± 5.6 and 29.0 ± 9.8 ms, respectively, for the 16 cells stimulated with the natural scene movie.

Events were determined as follows. The first spikes of bursts were extracted (Fig 1E, top) and merged across different repeats of the stimulus presentation (black in Fig 1E, middle). The intervals of these merged first spikes were then measured and a histogram was generated (Fig 1F). The histogram had two peaks separated by *T*_*thresh*_ (red and blue in Fig 1F), indicating that the intervals were composed of short intervals representing inter-trial fluctuation (red in Fig 1E, middle, and red in Fig 1F) and longer intervals separating the consecutive events (blue in Fig 1E, middle, and blue in Fig 1F). Thus, if two consecutively merged first spikes were closer than *T*_*thresh*_, they were incorporated into the same event; otherwise, they were assigned into two consecutive events (Fig 1E, bottom). The robustness of the method was evaluated as follows. If bursts occurred with a large timing jitter in different repeats, two consecutive bursts in one repeat were incorporated into one event. However, this occurred for only 2.7% ± 2.7% of bursts for the cells (mean ± SD, *n* = 41) stimulated with the spatially uniform stimulation, and 3.5% ± 1.1% for the cells stimulated with the natural scene movie (*n* = 16). The values were small, indicating that the definition of events was robust.

### Experimental Design and Statistical Analyses

The numbers of the analyzed cells and retinae were as follows. Salamanders: *n* = 41 cells in 15 retinae for spatially uniform stimulation, and *n* = 16 cells in 4 retinae for natural scene stimulation.

Correlations between the spike number and the intraburst ISIs of the bursts were investigated using data from ganglion cells stimulated with the spatially uniform stimulation as follows. For each event *j* in which at least one 2-spike burst occurred, the average number of spikes (*n*^(*j*)^) and the average intraburst ISIs of the 2-spike bursts (*m*^(*j*)^) were determined. The correlation between *n*^(*j*)^ and *m*^(*j*)^ was then calculated across events. Similar correlations were calculated for ISI_1_ and ISI_2_ of the 3-spike bursts.

Mutual information conveyed by the burst spike number was determined as follows. When the stimulation was repeated, bursts occurred in a limited number of discrete events that were defined at specific time points of the sequence (Fig 1A). Therefore, it was investigated how far the receiver of bursts can specify the events by knowing the spike number, as compared to the case where the receiver receives bursts without knowing the spike number. *N*_Rep_ and *N*_Ev_ represent the number of stimulus repeats and the number of events, respectively. 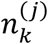 represents the number of *k*-spike bursts that occurred in the *j*-th event during the *N*_Rep_ repeats of stimulation (*k* = 0,1, 2, …, *k*_max_; *j* = 1, …, *N*_Ev_), where *k*_max_ is the largest number of spikes in a burst. For all *j*, 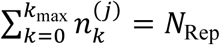. The number of all bursts that occurred in the *j*-th event is 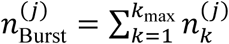, and the total number of bursts is 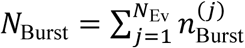. When a burst was generated, the probability that the burst was in the *j*-th event is 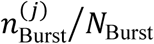. Thus, the prior entropy of events, calculated for the event probability after receiving a burst without knowing the spike number, is 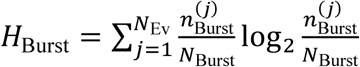. If the burst was *k*-spike, the posterior probability that the burst was in the *j*-th event is 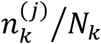, where 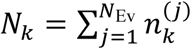 is the total number of *k*-spike bursts. Thus, the posterior entropy is 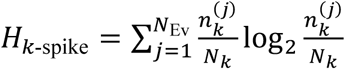. The mutual information is 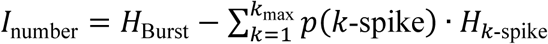, where *p*(*k*-spike) = *N*_3_/*N*_Burst_ is the probability of *k*-spike bursts. To evaluate the statistical significance, surrogate data were generated by exchanging the spike number of each burst with that of a randomly selected burst. One hundred surrogate data were generated and 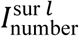 was calculated for the *l*-th surrogate (*l* = 1, …, 100). If *N*_larger_ of 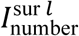 was larger than *I*_number_, the statistical significance is *P* = *N*_larger_/100. The estimated mutual information was corrected by the bias correction method [52].

Information conveyed by intraburst ISIs was calculated as follows. Intraburst ISIs of 2-spike bursts, and ISI_1_ and ISI_2_ of 3-spike bursts were divided into 4 groups according to the length of the ISIs, so that each group had as equal number of ISIs as much as possible. Two- and 3-spike bursts were thus divided into 4 and 16 groups, respectively. 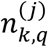 denotes the number of *k*-spike bursts of the group *q* in the *j*-th event (*q* = 1, …, *q*_max_; *q*_max_ = 4 for *k* = 2, *q*_max_ = 16 for *k* = 3). The total number of *k*-spike bursts of the group *q* is 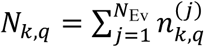. When a *k*-spike burst of the group *p* occurred, the posterior entropy is 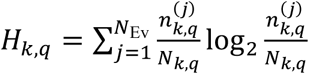. The information conveyed by ISIs is 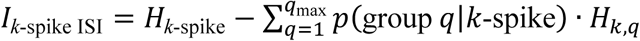, where *p*(group *q*|*k*-spike) = *N*_*k,q*_/*N*_*k*_. Statistical significance was evaluated similar to the spike number analysis. The information value was corrected for the bias [52].

Coordinate transformation of 3-spike burst ISIs was performed as follows (Fig 4B–D). For 3-spike bursts in event *j*, the average and standard deviation of ISI_1_ were designated as 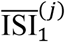 and 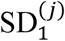, respectively. 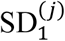 was linearly fitted with 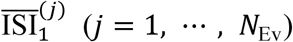 as 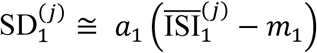, where *a*_1_ and *m*_1_ are constants (Fig 4C, top). For a variable 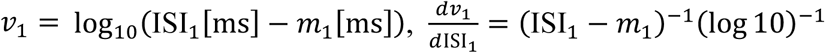, and thus the standard deviation of *v*_1_ in each event was similar among different events (see bursts shown in different colors in Fig 4D). Bursts with ISI_1_ ≤ *m*_1_ were removed from the analysis. *v*_2_ was similarly defined for ISI_2_ (Fig 4C, bottom). The principle axes of the distribution of *v*_1_ and *v*_2_ were determined (Fig 4D), and new variables *u*_1_ and *u*_2_ were defined by scaling *v*_1_ and *v*_2_ along these axes so that the standard deviations along these axes were equal to 1 (Fig 4E). *u*_1_ and *u*_2_ were shifted so that their averages were zero. The burst phase was atan2(*u*_2_, *u*_1_) (Fig 4E).

To characterize the stimulus features encoded by bursts, the stimulus sequences preceding bursts of a specific spike number, intraburst ISIs within a specific range, or burst phase within a specific range, were collected. The average and SEM values of the collected sequences were then used for the analyses. Neurons that generated only small numbers of 2- or 3-spike bursts were removed from the analyses (see legends for Figs 3G, 5D, 6C, and 6D). For linear reconstruction, stimulus sequences preceding the spikes in three-spike bursts were collected and averaged. The average sequence was divided by three and then used as the STA for the reconstruction (Fig 5G– I).

## Data Availability

The central data and computer codes used in this paper are available the open science framework database at: https://osf.io/29ect/. Other data and codes are available upon request.

## Acknowledgements

We thank Dr. Markus Meister for providing us with unpublished data and unstinting encouragement. We also thank Takao K. Hensch, Shun-ichi Amari, Shin Yanagihara, Neal Hessler, Yoshihiro Yoshihara, Stephan A. Baccus, Bence P. Ölveczky, Nick Lesica, Xin Jing, Charles Yokoyama, Naoki Masuda, Shiro Ikeda, Makoto Kaneda, Tomoki Fukai, Keita Watanabe, and Tetsuya Haga for thoughtful discussions.

## Author contributions

**Conceptualization**: Toshihiko Hosoya.

**Data curation**: Toshiyuki Ishii, Toshihiko Hosoya.

**Formal analysis**: Toshiyuki Ishii, Toshihiko Hosoya.

**Funding acquisition**: Toshihiko Hosoya.

**Investigation**: Toshiyuki Ishii.

**Methodology**: Toshiyuki Ishii, Toshihiko Hosoya.

**Project administration**: Toshihiko Hosoya.

**Resources**: Toshihiko Hosoya.

**Software**: Toshihiko Hosoya.

**Supervision**: Toshihiko Hosoya.

**Validation**: Toshiyuki Ishii, Toshihiko Hosoya.

**Visualization**: Toshiyuki Ishii, Toshihiko Hosoya.

**Writing – original draft**: Toshihiko Hosoya.

**Writing – review & editing**: Toshiyuki Ishii, Toshihiko Hosoya.

